# Loss of Ing4 enhances hematopoietic regeneration in multipotent progenitor cells

**DOI:** 10.1101/2024.09.27.615522

**Authors:** Georgina A. Anderson, Marco Hernandez, Carlos Alfaro Quinde, Zanshé Thompson, Melanie Rodriguez, Vera Binder-Blaser, Alison M. Taylor, Katie L. Kathrein

## Abstract

Despite its critical role in survival, many aspects of hematopoiesis remain unresolved. In the classical model of the hematopoietic program, quiescent hematopoietic stem cells (HSCs) sit at the top of the hematopoietic hierarchy, with the ability to self-renew and differentiate as needed. HSCs give rise to more proliferative progenitor cells, which possess multipotent potential, but have largely or completely lost self-renewal capabilities. Here, we have identified the tumor suppressor, Inhibitor of Growth 4 (Ing4), as a critical regulator of multipotent progenitor (MPP) homeostasis. In the absence of Ing4, we show that MPPs express a transcriptional program of hematopoietic activation, yet they remain quiescent with low levels of reactive oxygen species. Functionally, Ing4-deficient MPPs are capable of robust regeneration following competitive bone marrow transplantation, resulting in substantially higher blood chimerism compared to wild-type MPPs. These data suggest Ing4 deficiency promotes a poised state in MPPs, quiescent, but transcriptionally primed for activation, and capable of converting the poised state into robust repopulation upon stress. Our model provides key tools for further identification and characterization of pathways that control quiescence and regeneration in MPPs.

## Background

The diverse cells of the immune system are produced through the process of hematopoiesis.^1^ Stem cells capable of long-term donor engraftment are often described as long-term hematopoietic stem cells (LT-HSCs).^2,3^ LT-HSCs transition to ST-HSCs, which mature to multipotent progenitor cells (MPPs).^4,5^ The term “multipotent progenitor” is a general descriptor of the subset of heterogeneous HSPCs from the bone marrow (BM) that typically have lost self-renewal capability, do not contribute to serial transplantation, and are metabolically active.^6–9^

Several recent studies have uncovered a significant contribution of MPPs to hematopoiesis. These studies reveal that MPPs can supply long-term hematopoietic program and function during hematopoietic recovery without the contribution of HSCs.^7,10,11,13,15,17,19^ This body of work highlights the importance of the MPP population, but the field remains in the early stages of understanding the potential significance of these cells to hematopoiesis, particularly with regards to the mechanistic understanding of their contribution.

Ing4 is a tumor suppressor protein, generally localized to the nucleus, that is associated with a high frequency of acquired, inactivating mutations and poor prognoses in diverse human cancers.^12,14,16,18^ Ing4 has many regulatory roles, both as a chromatin remodeling protein within the Hbo1 complex and as a direct regulator of several major signaling pathways: NF-KB, c-Myc, p53, and HIF-1α.^20–25^ We have recently identified that Ing4-deficient HSCs are markedly different from wild-type HSCs, and maintain a quiescent, yet poised state ^26^. Here, we extend this line of inquiry by characterizing multipotent progenitor cells from the Ing4^-/-^ mouse model. In the Ing4^-/-^ bone marrow, MPPs are less abundant that wild-type (WT) MPPs and express an activation profile. Yet, Ing4^-/-^ MPPs are more quiescent than WT MPPs and do not convert the activation profile into functional activation at steady-state. Upon stress, however, Ing4^-/-^ MPPs show significantly increased engraftment in sorted competitive BM transplantation compared to WT MPPs.

## Results

### MPP Differentiation is impaired in the absence of Ing4

To profile multipotent progenitor cells in the absence of Ing4, we used a previously described Ing4-deficient mouse model.^21^ Multipotent progenitors are defined here as the Lin^-^ Sca1^+^ cKit^+^ CD48^-^ CD34^+^ CD150^-^ fraction of the whole bone marrow (Gating strategy in **Figure 1A**). Cell surface profiling of steady-state bone marrow showed a significant decrease in MPPs as a percentage of LSKs observed in BM from Ing4^-/-^ mice (Figure 1 A,B). This may potentially result from a differentiation block at the ST-HSC stage.^26^ No difference in cell size is observed between WT and Ing4^-/-^ MPPs when comparing MFIs of FCS profiles (Figure 1C).

**Figure 1.**
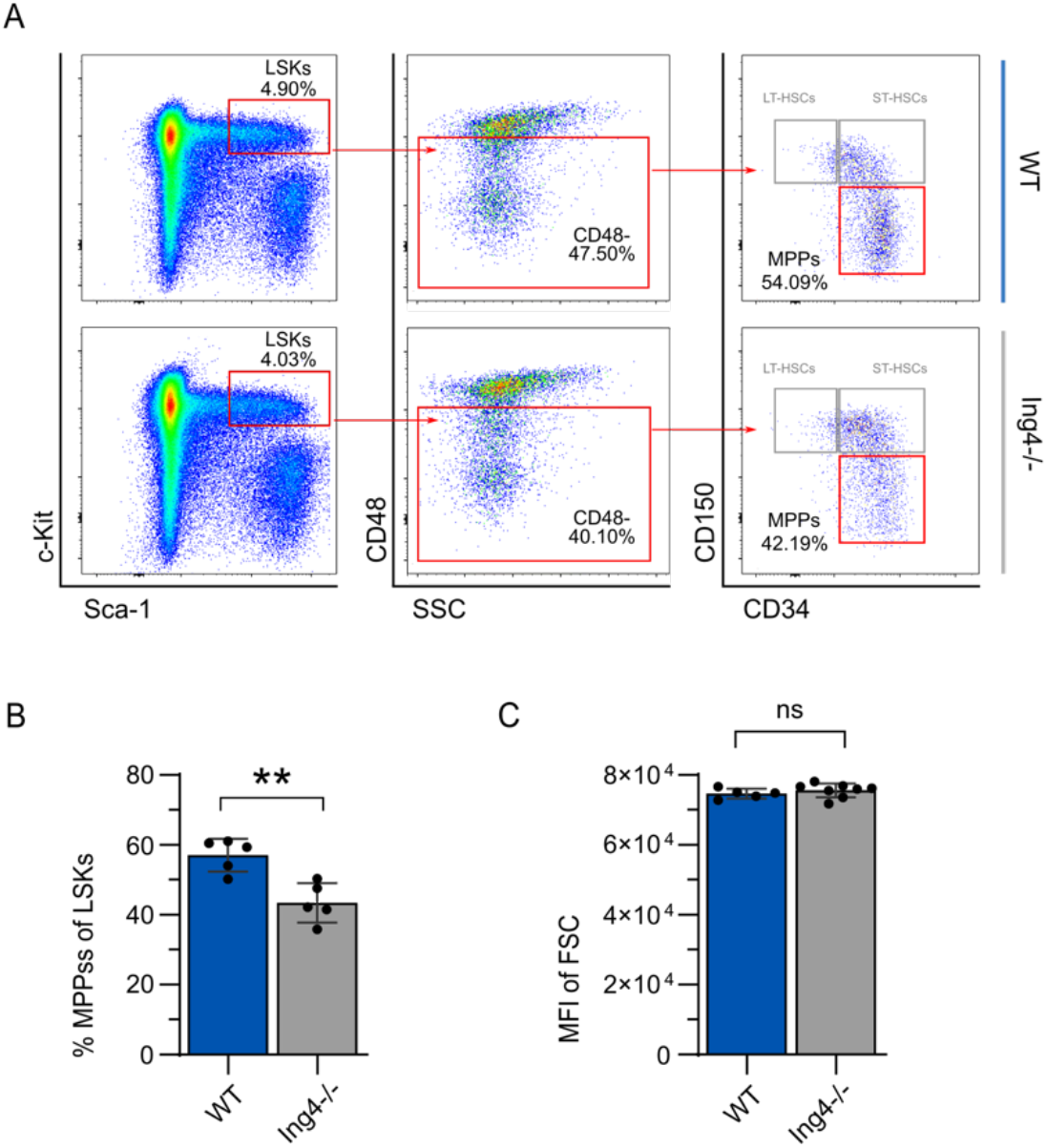
Steady state MPP populations from WBM of wildtype and Ing4^-/-^ mice. (A) Representative gating strategy of flow cytometric analysis for MPPs. (B) MPPs from WBM isolated from individual WT and Ing4^-/-^ steady state mice as a percentage of LSK cells. (*n*=10; \*\**p*<0.01). (C) Flow cytometric analysis of cell size by FSC of MPPs isolated from individual WT and Ing4^-/-^ mice (*n*=5-8; *=*p*<0.05). Statistical significance was assessed using Student’s t-test with Welch’s correction.

### Ing4-deficient MPPs show characteristics of quiescence

Quiescence is a state in which the cell has the ability to divide, but largely remains in a resting state under steady-state conditions.^27^ Cells can enter the quiescent, dormant G_0_ phase from G_1_, thereby minimizing susceptibility to mutations in the DNA acquired through cell division and maintain long-term preservation of hematopoiesis.^28,29^ MPPs are more metabolically active, and spend more time in G_1_ than HSCs^7^.

To examine cell cycle status in the absence of Ing4, 4’-6-diamidino-2-phenylindole (DAPI) was used to visualize DNA content and Ki-67 showed proliferative status. When compared to WT, cell cycle status of Ing4^-/-^ MPPs showed an increased proportion of cells in G_0_ (**Figure 2A and B**).

**Figure 2.**
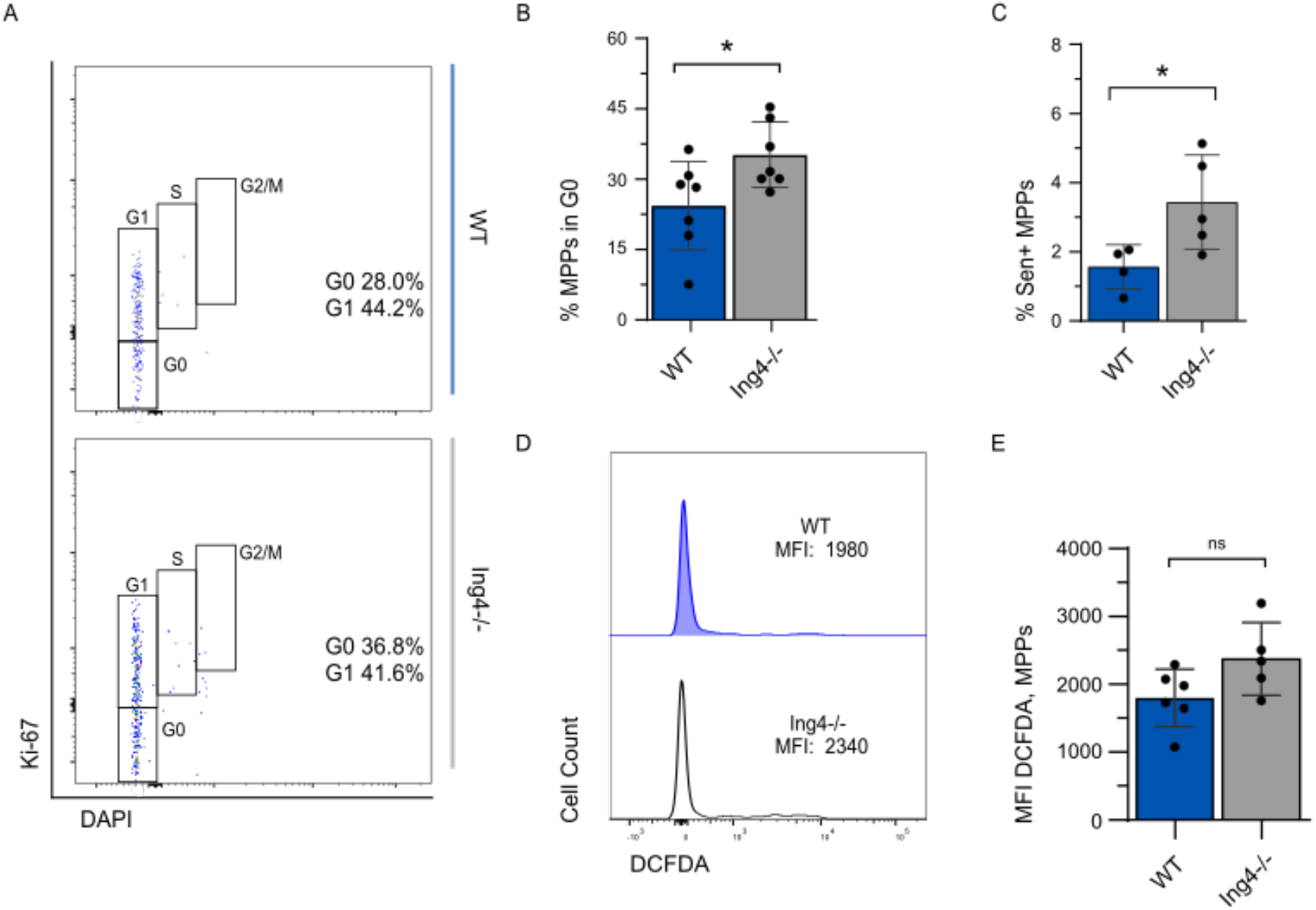
Ing4-deficient MPPs have increased quiescence and low ROS levels. (A) Representative flow cytometric analysis of MPPs using Ki-67 and DAPI for cell cycle profile. (B) MPPs in G_0_ isolated from WBM of individual WT and Ing4^-/-^ steady state mice as a percentage of the MPP population. (*n*=4-7; *=*p*<0.05). (C) Representative flow cytometric analysis of beta-galactosidase staining for senescence in MPP populations in WT and Ing4^-/-^ bone marrow at steady state. (D) Senescence-positive MPPs isolated individual WT and Ing4^-/-^ steady state mice as a percentage of all MPPs. (*n*=4-5; *=*p*<0.05). (E) Representative flow cytometric analysis of staining with DCFDA for ROS in MPPs from BM of WT and Ing4^-/-^ mice. (F) MFI of DCFDA in MPPs isolated from WT and Ing4^-/-^ steady state mice. (*n*=5-6; ns=*p*>0.05). Statistical significance was assessed using Student’s t-test with Welch’s correction.

Senescence is defined as irreversible stable cell cycle arrest and happens in response to intrinsic or extrinsic stressors. Because Ki-67 only differentiates between cycling versus non-cycling cells, it cannot discriminate between quiescent and senescent cells, as cells in neither state will incorporate Ki-67. Analysis of senescence-associated β-galactosidase (SA-β-gal) staining of WT and Ing4^-/-^ MPPs revealed a small but significant increase in percentage of senescence-positive MPPs (1.6% and 3.4%, respectively) (**Figure 2C and D**). The relatively small proportions of senescent-positive cells found here likely do not account for the difference in proportions of MPPs observed to be in the G_0_ phase of the cell cycle, suggesting that MPPs from Ing4^-/-^ mice are more quiescent than WT.

As low intracellular reactive oxygen species (ROS) content is associated with cell quiescence,^30,31^ MPPs were assayed to determine if ROS levels differed between WT and Ing4^-/-^ MPPs. Lineage-depleted BM cells were treated with 2’7’-deichlorofluorescein diacetate (H_2_DCFDA) to examine ROS levels in WT and Ing4^-/-^ MPPs. ROS levels, as indicated by mean fluorescence index, showed no significant difference between WT and Ing4^-/-^ MPPs (**Figure 2E and F**).

To investigate if loss of Ing4 impacts apoptosis in MPPs and may account for the reduction in MPPs overserved in the absence of Ing4, we determined the frequency of annexin V^+^ MPPs undergoing apoptosis in WT and Ing4-deficient populations. The percentage of MPPs undergoing apoptosis was similar between WT and Ing4^-/-^ mice (**Supplemental Figure 1**).

### Ing4^-/-^ MPPs Simultaneously Express Genes Associated with Activation and Quiescence

To elucidate the molecular consequences of Ing4 loss in HSC regulation, we conducted a genome-wide expression analysis using bulk RNA-sequencing (RNA-seq) of purified Ing4^-/-^ LT-HSCs (LSK CD48^-^CD34^-^CD150^+^), ST-HSCs (LSK CD48^-^CD34^+^CD150^+^) and MPPs. We then compared the profiles of these populations, pooled, against pooled wild-type LT-HSC, ST-HSC, and MPP gene expression. This analysis provided a way to identify stem cell-like gene signatures in the MPP populations. We revealed 1,635 differentially expressed genes in the combined HSPC populations (936 upregulated and 699 downregulated, p<0.05). Surprisingly, Gene Set Enrichment Analysis (GSEA) of this data set using the Hallmark gene set showed common upregulated genes in Ing4^-/-^ HSPCs were associated with oxidative phosphorylation, ribosomal biogenesis (RiBi), and c-Myc target gene expression (**Figure 3A**). Downregulated genes were associated with mitotic spindle formation, and UV response (**Figure 3A**). All dysregulated genes are represented in a volcano plot shown in **Figure 3B**. A set of genes that are associated with quiescence and cell cycle regulation were also upregulated, including the cell-cycle regulator *p57, txn1* and *gpx1* (**Figure 3B**).

**Figure 3.**
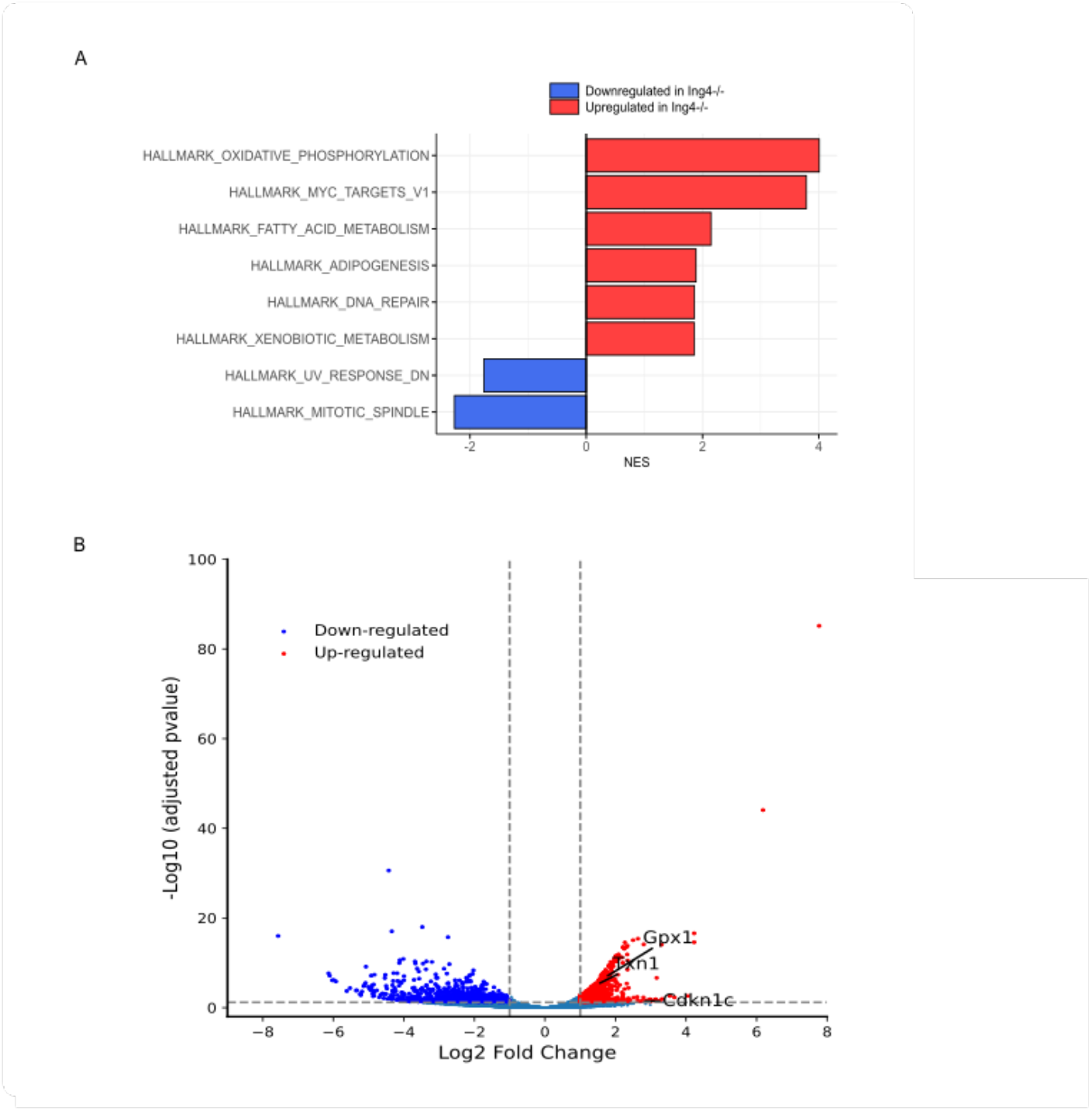
Loss of Ing4 promotes expression of a hematopoietic activation profile in MPPs. (A) Normalized Enrichment Scores (NES) from a Hallmark gene set analysis comparing populations (WT LT-HSC, ST-HSCs, and MPP pooled), and (Null LT-HSC, ST-HSCs, and MPPs pooled). A total of 8 enrichment pathways (FDR-adjusted *p*-value<0.05) were identified. Bar colors indicate statistical significance. (B) Differentially expressed genes in pooled WT HSPCs compared to pooled Ing4^-/-^ MPPs, with a statistically significant threshold of *p*<0.05, and a Log_2_Fold change =1. Upregulated genes are represented in red, downregulated in blue.

### Ing4^-/-^ MPPs have normal levels of translation and OXPHOS utilization

Based on the RNA-sequencing data, we next sought to determine if Ing4^-/-^ MPPs are more metabolically active, as their transcriptional profile would suggest. Hematopoietic activation is associated with increased translation (Signer et al., 2014).To examine if an increase in translation rate accompanies the increase in RiBi asocated genes upregulated in our RNA-seq data set, we used O-propargyl-puromycin (OP-Puro) incorporation. To this end, lineage depleted bone marrow cells were treated with OP-Puro for 1 hour, then analyzed for levels of OP-Puro incorporation. Surprisingly, these assays show no increase in the translation rates of Ing4^-/-^ MPPs (**Figure 4A**).

**Figure 4.**
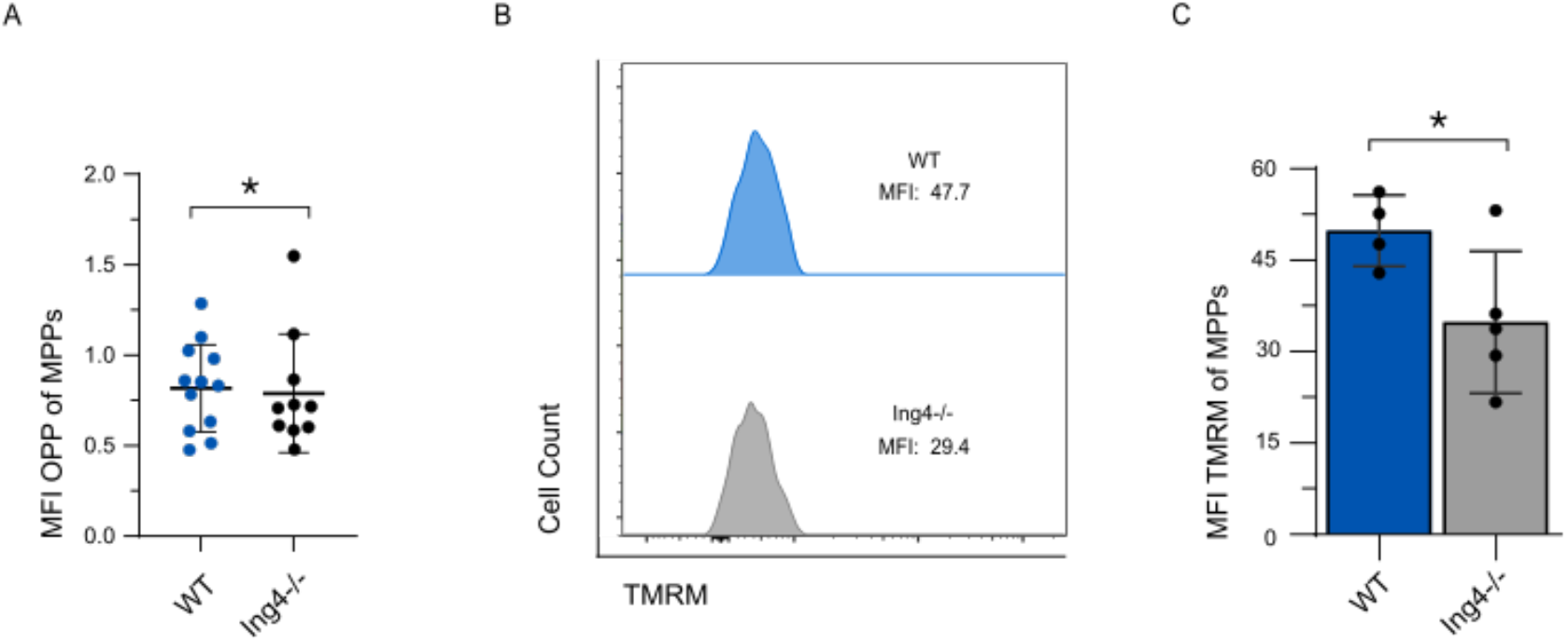
Ing4-deficient MPPs have normal translation rates and reduced mitochondrial potential. (A) MFI of OPP-treated MPPs isolated from individual WT and Ing4^-/-^ steady state mice. (*n*=10-23; * *p*<0.05). (B) Representative flow cytometric analysis of staining for TMRM in MPP populations in WT and Ing4^-/-^ WBM at steady state. (C) MFI of TMRM-treated MPPs isolated from individual WT and Ing4^-/-^ steady state mice. (*n*=5; * *p*<0.05). Statistical significance was assessed using Student’s t-test with Welch’s correction for (A) and Mann-Whitney analysis (B).

To further address Ing4^-/-^ MPP activation, we examined the OXPHOS pathway, as components of this pathway also show transcriptional activation. As HSPCs are activated, there is a characteristic shift from glycolysis to oxidative phosphorylation. To profile metabolic activation, we examined tetramethylrhodamine methyl ester (TMRM) levels which indicate mitochondrial membrane potential. As activated cells transition from glycolysis to oxidative phosphorylation to generate ATP, an electrical potential gradient forms across the mitochondrial membrane and is observable in TMRM staining. We observed a significant decrease in MFI in Ing4^-/-^ MPPs as compared to WT counterparts, suggesting that Ing4^-/-^ MPPs are not using OXPHOS to generate higher levels of ATP than WT MPPs (**Figure 4B, C**). The delocalization of TMRM in the mitochondria indicated by the decreased MFI in the Ing4^-/-^ MPPs suggests they may rely more heavily on glycolysis to generate ATP than WT MPPs.

Together, these signatures suggest that Ing4 deficiency may promote a transcriptionally poised state in MPPs, whereby Ing4 loss upregulates genes associated with activation while also inducing quiescence-associated genes to maintain MPPs in a resting state.

### Ing4^-/-^ MPPs contribute to multilineage engraftment at higher levels than WT MPPs

To compare the engrafting capacities of Ing4^-/-^ and WT MPPs, we sorted MPPs from CD45.2+ Ing4^-/-^ and WT donor mice. Separately, pools of either 100 Ing4^-/-^ or 100 WT MPPs were transplanted, along with 200,000 CD45.1+ competitor marrow cells into CD45.1+ recipient mice (**Figure 5A**). Peripheral blood was analyzed at 12 weeks post-transplant and revealed a significant increase of donor contribution from the Ing4^-/-^ MPPs than WT (61.3% and 18.6%, respectively) (**Figure 5B**). Furthermore, Ing4^-/-^ MPPs were shown to reconstitute the myeloid and T-cell populations at a significantly higher rate than WT MPPs (**Supplemental Figure 2**). Transplanted Ing4^-/-^ MPPs also made a greater contribution to B-Cell population than WT MPPs, though this difference was not statistically significant. Moreover, this increase in chimerism is observed up to 9 months post-transplantatoin (**Figure 5C**), providing clear evidence for the unexpectedly robust engraftment potential of MPPs from Ing4^-/-^ donors.

**Figure 5.**
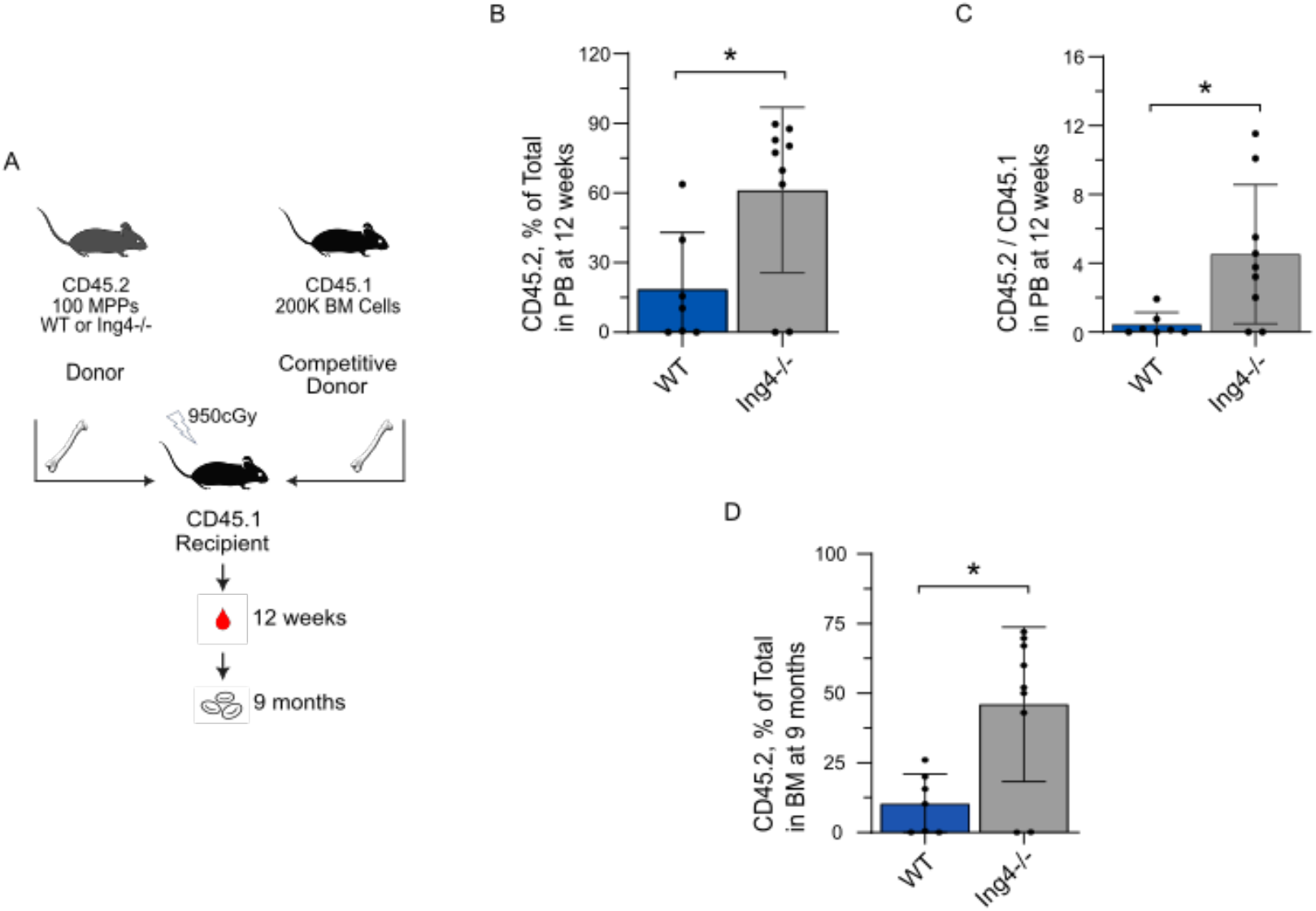
In a sorted, competitive bone marrow transplantation assay, Ing4^-/-^ MPPs contribute to chimerism at higher levels than WT MPPs. (A) Schematic overview of the sorted, competitive transplantation assay using BM from WT and Ing4^-/-^ mice. (B) CD45.2 chimerism in individual MPP-recipient mice from peripheral blood collected 12 weeks following sorted, competitive transplantation from WT or Ing4^-/-^ mice. (*n*=7-9; *=*p*<0.05). (C) CD45.2 chimerism in individual MPP-recipient mice from peripheral blood collected 9 months following sorted, competitive transplantation from WT or Ing4^-/-^ mice. (*n*=7-9; *=*p*<0.05). Statistical significance was assessed using Mann-Whitney analysis.

## Discussion

The mechanisms that regulate maintenance and proliferation of hematopoietic homeostasis in the bone marrow, including quiescence, self-renewal, and activation, remain elusive. Ing4 deficiency results a decreased proportion of MPPs from the LSK compartment as compared to WT MPPs, providing evidence of skewed hematopoiesis. MPPs from Ing4^-/-^ mice have increased quiescence over WT counterparts, with a much larger increase in non-cycling cells than can be accounted for by SA-β-gal^+^ senescent population. Strikingly, Ing4-deficient MPPs display much higher levels of engraftment than WT MPPs from Ing4^-/-^ donors and are able to support multilineage regeneration of the hematopoietic system.

Few models of enhanced MPP activity have been previously identified. Introduction of NUP98-HOXA10hd into MPPs resulted in long-term reconstitution up to 44-weeks post-transplant, similar to our observations with Ing4^-/-^ MPPs. Similarly, ectopic Sox17 expression in adult HSCs and MPPs also confers increased self-renewal potentiall, where HSPCs are more reminiscent of fetal HSCs.^33^ Sox17 expression in adult HSPCs allowed for long-term multilineage reconstitution of the hematopoietic system following transplantation, though skewed toward myelopoiesis and erythropoiesis, which, again, bears resemblance to fetal hematopoiesis patterns.^33^ Finally, ectopic expression of miR-125a in murine and human multipotent progenitors also resulted in enhanced self-renewal and long-term multilineage engraftment following serial transplantation.^34–37^ Although the pathways altered in these different models are largely varied amongst each other and our model, taken together, these data provide strong evidence of the potential for MPPs to have an impact on enhancing hematopoietic recovery.

Recent work in characterization of MPP4 and 5 subpopulations suggest MPPs can retain an increased capacity for quiescence and differentiation, and further highlight the contributions of MPP subpopulations to the maintenance of the hematopoietic system under steady-state conditions. ^7,38^ Together, these data show that MPPs with the capacity to regenerate at substantial levels may have clinical significance and warrant continued study.

### Limitations of Study

MPPs are described by the presence of a wide variety of cell surface markers that can be profiled by flow cytometry. While we have defined MPPs as Lin-Sca-1^+^ c-Kit^+^ CD150^-^ CD34^+^, we have not used CD135, CD244 or CD229, which are additional commonly used cell surface markers in isolating subpopulations of HSPCs. This, coupled with deeper analysis of RNA from Ing4^-/-^ MPPs would likely provide insight into molecular pathways that may confer this cell population with its unique characteristics.

## Methods

### Ing4^-/-^ Mouse Model

Ing4^-/-^ mice (CD45.2) were provided by Stephen N. Jones (University of Massachusetts Medical School) and colony was maintained on site. Wildtype (WT) C57BL/6 (CD45.2) and SJL-Ptprc^a^ Pepc^b^/BoyJ (CD45.1) mice 8-12 weeks of age served as controls. CD45.1 mice were purchased from The Jackson Laboratory. All mice were maintained at an AALAC-accredited animal facility at the University of South Carolina according to Institutional Animal Care and Use Committee animal guidelines, and all animal experiments were performed with consent from the local ethical committee.

### Bone Marrow Collection and MPP Isolation

The Lineage^-^ Sca^+^ c-Kit^+^ (LSK) CD48^-^ CD34^+^ CD150^-^ cell fraction was isolated from WBM. Mice were humanely euthanized, and the long bones (femur, tibia, humerus, radius) were removed. Long bones from individual mice were crushed then filtered through a 40μm nylon cell strainer. Cells were incubated with 1x RBC Lysis Solution (Miltenyi Biotec) to lyse red blood cells. Cells were lineage depleted via a Lineage Cell Depletion Kit (Miltenyi Biotec) using LS columns and an LS magnetic separator.

### Collection of Peripheral Blood

Under anesthesia, peripheral blood was collected from retroorbital venous sinus into heparinized (0.05 IU/mL; Alfa Aesar) micro-hematocrit capillary tubes (Kimble). Cells were incubated with 1x RBC Lysis Solution (Miltenyi Biotec) to lyse red blood cells.

### Antibody Staining

After lineage depletion, BM cells were incubated for in the dark for 30 minutes at 4°C with antibodies for MPP profiling (Lineage: B220, CD3e, CD11b, Gr-1, Ter119; Sca-1, c-Kit, CD48, CD34, CD150). Cells were then rinsed twice with phosphate-buffered saline (PBS)-bovine serum albumin (BSA). Antibodies against CD45.2 and CD45.1 were used to assess engraftment following transplantation assays. See Table 2.2 for complete list of antibodies and concentrations.

### Reactive Oxygen Species Assay

Lineage-depleted BM cells were stained for MPP profiling, then incubated with 7.5 μM H_2_DCFDA (Invitrogen) in PBS for 30 minutes at 37°C. Cells were rinsed with PBS and analyzed by flow cytometry.

### Cell Fixing and Permeabilization

Lineage-depleted, stained WBM cells were fixed and permeabilized with Cytofix/Cytoperm kit (Invitrogen) according to manufacturer’s instructions.

### Cell Cycle Characterization

Fixed and permeabilized cells were incubated in the dark overnight at 4°C with Ki-67 antibody (Invitrogen), then rinsed with PBS. Prior to analysis via flow cytometry, cells were incubated for 15 minutes at room temperature with DAPI (0.2 μL/mL, Invitrogen) and rinsed with PBS.

### Senescence Assay

Lineage-depleted, stained BM cells were fixed and permeabilized with Cytofix/Cytoperm kit (Invitrogen) according to manufacturer’s instructions. Fixed and permeabilized cells were incubated with CellEvent Senescence Probe (Thermo) for 2 hours at 37ºC. Cells were rinsed with PBS. Cells were analyzed by flow cytometry.

### Sorted Bone Marrow Transplant

To compare the reconstitution and maintenance capacity of Ing4-deficient MPPs, sorted MPPs from CD45.2 Ing4^-/-^ and WT mice were transplanted in a competitive setting. Either 100 Ing4^-/-^ or 100 WT MPPs were combined with 200,000 CD45.1 unfractionated WBM cells and, under anesthesia, transplanted via retroorbital injection into 8-to 12-week-old recipient mice (B6. SJL-CD45.1) that had been lethally irradiated (9.5 Gy, administered as two doses at least three hours apart). Peripheral blood and BM of recipient mice were analyzed at 12 weeks post-transplant.

### OP-Puromycin staining

Lineage-depleted BM cells were stained for MPP profiling, then incubated with preheated DMEM solution containing 25uM OP-Puromycin (Invitrogen) for 60 minutes as described previously^39^. Cells were rinsed with PBS and the Click-iT™ azide-alkyne cycloaddition reaction was conducted per manufacturers protocol, then cells were analyzed by flow cytometry.

### Flow Cytometry, Cell Sorting and Analysis

Cells were analyzed on LSR II (BD), FACSAria II (BD), or FACSymphony (BD). For sorted cell assays, FACSAria II (BD) was used. FACS Diva was used in conjunction with flow cytometers for acquisition and FlowJo (version 10) was used for analysis of MPP populations.

### Statistical Analysis

Mean values ± SD are shown. Student’s t test with Welch’s correction or Mann-Whitney analysis were used for single comparisons (GraphPad Prism v.10.1.1). \**p*<0.05, \*\**p*<0.01, \*\*\**p*<0.005, \*\*\*\**p*<0.001. Two-way ANOVA was used for multiple comparisons for in vivo inhibitor assays (GraphPad Prism v.10.1.1).

## Figure Legends

**Supplemental Figure 1.**
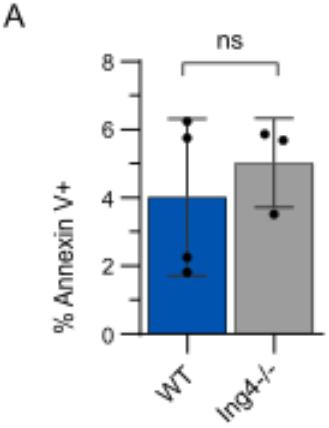
MPPs from Ing4^-/-^ and WT mice undergo similar rates of apoptosis. Percentage of Annexin V^+^ MPPs of individual WT and Ing4^-/-^ steady state mice. (*n*=3-4; ns=*p*>0.05). Data in S1 reflect mean values ± SD. Statistical significance was assessed using Mann-Whitney analysis.

**Supplemental Figure 2.**
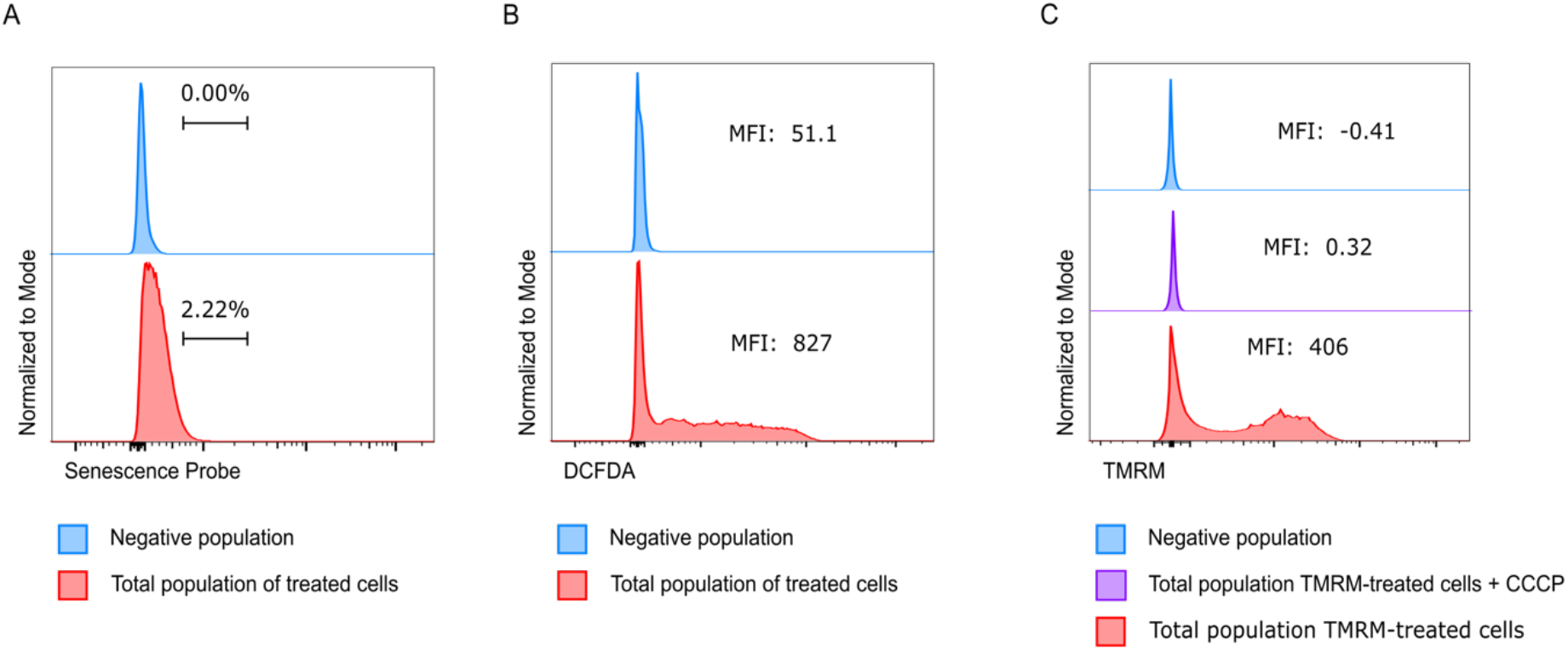
Control staining for senescence, reactive oxygen species, and mitochrondrial membrane potential in control samples. (A) Representative flow cytometric analysis of unstained (blue) and all cells (red) stained with beta-galactosidase for senescence. (B) Representative flow cytometric analysis of unstained (blue) and all cells (red) stained with DCFDA. (C) Representative flow cytometric analysis of unstained (blue), all cells treated with CCCP and stained with TMRM (purple) and all cells stained with TMRM (red) for mitochondrial potential. (*n*=4-7).

**Supplemental Figure 3.**
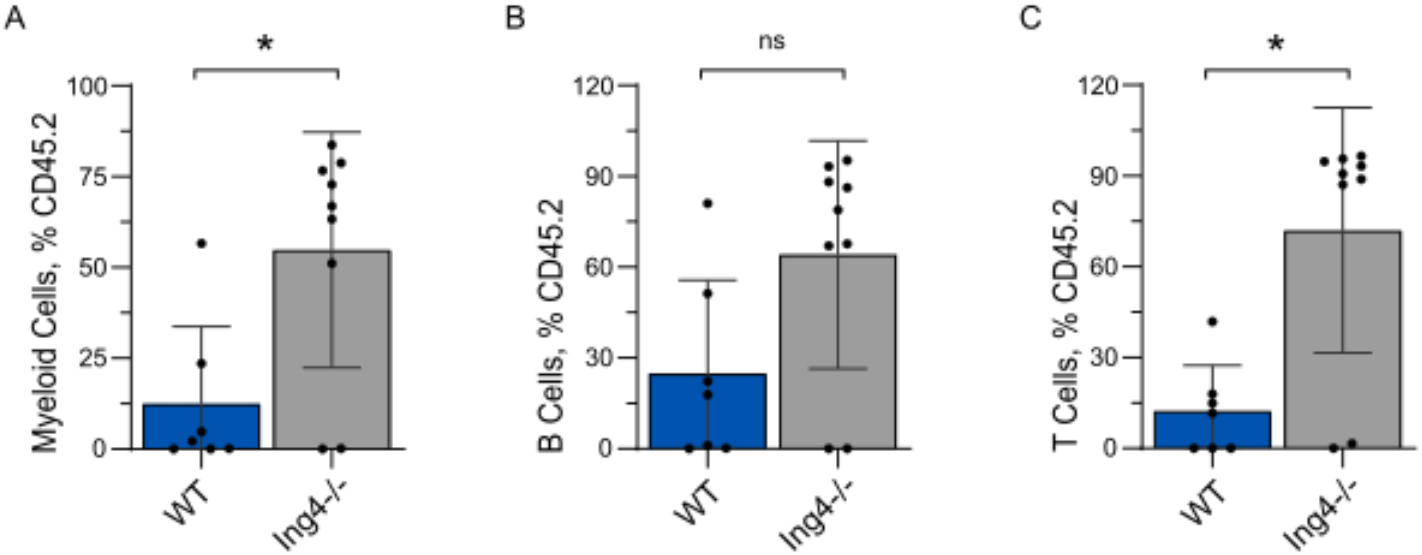
Bone Marrow chimerism of individual lineages shows mild lineage bias towards Myeloid and T cells. (A) CD45.2 chimerism of myeloid cells in individual MPP-recipient mice from peripheral blood collected 12 weeks following sorted, competitive BM transplant from WT or Ing4^-/-^ mice. (*n*=7-9; *=*p*<0.05). (B) CD45.2 chimerism of B cells in individual MPP-recipient mice from peripheral blood collected 12 weeks following sorted, competitive BM transplant from WT or Ing4^-/-^ mice. (*n*=7-9; ns=*p*>0.05). (C) CD45.2 chimerism of T cells in individual MPP-recipient mice from peripheral blood collected 12 weeks following sorted, competitive BM transplant from WT or Ing4^-/-^ mice. (*n*=7-9; *=*p*<0.05). Data reflect mean values ± SD. Statistical significance was assessed using Mann-Whitney analysis.

